# TAF7L REGULATES EARLY STAGES OF MALE GERM CELL DEVELOPMENT

**DOI:** 10.1101/2023.10.08.561408

**Authors:** Ayelen Moreno-Irusta, Esteban M. Dominguez, Khursheed Iqbal, Xiaoyu Zhang, Ning Wang, Michael J. Soares

## Abstract

Male germ cell development is dependent on the orchestrated regulation of gene networks. TATA-box binding protein associated factors (**TAFs**) facilitate interactions of TATA-binding protein with the TATA element, which is known to coordinate gene transcription during organogenesis. TAF7 like (***Taf7l***) is situated on the X chromosome and has been implicated in testis development. We examined the biology of TAF7L in testis development using the rat. *Taf7l* was prominently expressed in preleptotene to leptotene spermatocytes. To study the impact of TAF7L on the testis we generated a global *loss-of-function* rat model using CRISPR/*Cas9* genome editing. Exon 3 of the *Taf7l* gene was targeted. A founder was generated possessing a 110 bp deletion within the *Taf7l locus*, which resulted in a frameshift and the premature appearance of a stop codon. The mutation was effectively transmitted through the germline. Deficits in TAF7L did not adversely affect pregnancy or postnatal survival. However, the *Taf7l* disruption resulted in male infertility due to compromised testis development and failed sperm production. Mutant germ cells suffer meiotic arrest at the zygotene stage, with defects in sex body formation and meiotic sex chromosome inactivation. This testis phenotype was more pronounced than previously described for the subfertile *Taf7l* null mouse. We conclude that TAF7L is essential for male germ cell development in the rat.

## INTRODUCTION

Spermatogenesis is a highly organized differentiation process that relies on multiple factors. Precise temporal and spatial patterns of transcription factor expression and action drive the normal progression of spermatogenesis (1–3). Molecular events regulating spermatogenesis include germ cell-specific core transcriptional machinery and the involvement of the TATA-binding protein (**TBP**) family (2, 3). The RNA polymerease II general transcriptional factor (**TFIID**) complex contains members of the TBP family and TBP-associated factors (**TAFs**)(4, 5). TAFs have been directly linked to the regulation of spermatogenesis (6–10). TBP associated factor 7 like (**TAF7L**) is an X-linked homologue of TBP associated factor 7 (**TAF7**) expressed in the testis (11–13).

TAF7L has been implicated in the regulation of male gamete development in the mouse and human. In the mouse, TAF7L is expressed during spermatogenesis from the spermatogonia stage to the round spermatid stage. Mice deficient in TAF7L exhibit subfertility and abnormalities in sperm structure and motility (12, 13). TAF7L has also been linked to spermatogenesis and male fertility in the human (14–16). A missense mutation in the human *TAF7L* gene resulting in an aspartate to glycine amino acid change (D136G) has been identified as a potential cause of oligozoospermia (17), while other mutations within the *TAF7L* gene in men contributing sperm for in vitro fertilization have been linked to poor outcomes (18).

In this report, we assessed the role of TAF7L in rat spermatogenesis. We generated a *Taf7l* mutant rat using CRISPR/Cas9 genome editing. Disruption of *Taf7l* resulted in male infertility secondary to an arrest in meiosis. We also demonstrated the involvement of TAF7L in the regulation of meiotic sex chromosome inactivation (**MSCI**). These observations highlight differences in the biology of TAF7L in the mouse versus the rat testes and provide an alternative model for investigating the involvement of TAF7L in regulating spermatogenesis, which may have relevance to human male infertility.

## MATERIALS AND METHODS

### Animals and tissue collection

Holtzman Sprague-Dawley rats were maintained on 14:10 h (light:dark cycle), with free access to food and water. Males were euthanized between 2 to 12 weeks of age and testes collected. Some testes were frozen in dry ice-cooled heptane and stored at - 80°C, or fixed in Bouin’s solution and embedded in paraffin, for histological analyses, whereas other testes were dissected, weighed, and tissues frozen in liquid nitrogen and stored at -80°C until used for RNA analyses. The University of Kansas Medical Center Animal Care and Use Committee approved all protocols used in this report.

### Generation of a *Taf7l* mutant rat model

Mutations at the *Taf7l* locus were generated using CRISPR/Cas9 genome editing (19, 20). Guide RNAs targeting Exon 2 (target sequence: GTTCATATTGCGTCTGCCAC; nucleotides 5585-5605) and Exon 3 (target sequence: TGTTTCACTGCCTGCTAAGC; nucleotides 7720-7740) of the *Taf7l* gene (NM_033230.3) were electroporated into single-cell rat embryos using the NEPA21 electroporator (Nepa Gene Co Ltd, Ichikawa City, Japan). Electroporated embryos were transferred to oviducts of day 0.5 pseudopregnant rats. Offspring were screened for *Taf7l* mutations using REDExtract-N-Amp^TM^ Tissue PCR kit (XNAT, Millipore Sigma, Burlington, MA) extracted genomic DNA from tail-tip biopsies. Polymerase chain reaction (**PCR**) was performed on the purified DNA samples using primers flanking the guide RNA sites (Forward primer: GCTTATCTAGCATGCGCAAA, Reverse primer: GTAAAATACAATATGAAAAAGCAAGC). PCR products were resolved by agarose gel electrophoresis and identified using ethidium bromide staining. Genomic DNA samples containing potential mutations were amplified by PCR, gel purified, and precise boundaries of deletions determined by DNA sequencing (Genewiz Inc., South Plainfield, NJ). Founders with *Taf7l* mutations were backcrossed to wild type rats to demonstrate germline transmission. Only female founders were successful in transferring *Taf7l* mutations. Routine genotyping was performed by PCR on genomic DNA with primer sets presented above.

### Sperm quantification

Cauda epididymidis from wild-type (*Taf7l ^Xm+/Y^*) and mutant (*Taf7l^Xm−/Y^*) rats were dissected and separated from their associated fat pad, blood vessels and connective tissue. Sperm were collected by placing minced cauda epididymides in modified Tyrode’s medium (95 mM NaCl, 4.7 mM KCl, 1.2 mM KH_2_PO_4_, 1.2L mM MgSO_4_, 5.5 LmM glucose, 0.27 mM pyruvic acid, 0.25L mM lactic acid, 40L mM HEPES, and 20 LmM Tris, pH 7.4) (21) for 10Lmin at 37L°C. Sperm were counted using a hematocytometer. Sperm quantification was repeated four times for each sample.

### Testis histology

Testes from *Taf7l ^Xm+/Y^* and *Taf7l^Xm−/Y^* rats were fixed overnight in Bouin’s fixative at 4°C. Bouin’s fixed testes were washed three times in phosphate buffered saline (**PBS**, pH 7.4) for 10 min at 4°C, followed by three washes in 70% ethanol for 10 min at 4°C. Fixed tissues were stored in 70% ethanol prior to embedding in paraffin and sectioning (5 μm). Tissue sections were deparaffinized with xylene and rehydrated in a graded series of ethanol solutions and then transferred to PBS. Representative sections were stained with hematoxylin and eosin. TUNEL assays were performed using APO-Direct Kit (TNB-6611-R, Tonbo Biosciences, San Diego, CA) according to the manufacturer’s instructions. Images were captured on a Nikon 90i upright microscope with a Roper Photometrics CoolSNAP-ES monochrome camera.

### In situ hybridization

In situ detection of *Taf7l* and *Stra8* transcripts were performed on paraffin-embedded rat testis tissue using the RNAscope Fluorescent Multiplex Reagent Kit, version 2 (Advanced Cell Diagnostics, Newark, CA). Probes were prepared by Advanced Cell Diagnostics to detect rat *Taf7l* (860161, NM_001135877.1, target region: nucleotides 2 - 1044) and rat *Stra8* (1129161-C2, XM_006236282.4; target region: nucleotides 239 - 1372), Fluorescence images were acquired on a Nikon 90i upright microscope with a Roper Photometrics CoolSNAP-ES monochrome camera.

### RNA isolation and RT-qPCR

Total RNA was extracted using TRI Reagent Solution (AM9738, ThermoFisher, Waltham, MA) following the manufacturer’s instructions. Total RNA was reverse transcribed using a High-Capacity cDNA Reverse Transcription Kit (4368813, ThermoFisher) to generate testis cDNA, which was diluted 1:10 for RT-qPCR measurements using PowerUp SYBR Green Master Mix (A25742, ThermoFisher. QuantStudio 5 Flex Real-Time PCR System (Applied Biosystems, Foster City, CA) was used for amplification and fluorescence detection. RT-qPCR was performed under the following conditions: 95℃ for 10 min, followed by 40 cycles of 95℃ for 15 sec and 60℃ for 1 min. All reactions were performed in duplicate, and mean crossing point (**Cp**) values were used for the analysis. Relative mRNA expression was calculated using the delta-delta Ct method. Cp values were normalized to the value obtained for *Gapdh* reactions (ΔCp). Differences between *Taf7l ^Xm+/Y^* and *Taf7l^Xm−/Y^*samples were calculated for each primer set (ΔΔCp) and the fold change was calculated as 2^−ΔΔCp^.Primers used for RT-qPCR are provided in **Table S1**.

### RNA-seq analysis

Transcript profiles were generated from *Taf7l ^Xm+/Y^* and *Taf7l^Xm−/Y^*testes (PND 30 and 60). Complementary DNA libraries from total RNA samples were prepared with Illumina TruSeq RNA preparation kits according to the manufacturer’s instructions (Illumina, San Diego, CA). RNA integrity was assessed using an Agilent 2100 Bioanalyzer (Santa Clara, CA). Barcoded cDNA libraries were multiplexed onto a TruSeq paired-end flow cell and sequenced (100-bp paired-end reads) with a TruSeq 200-cycle SBS kit (Illumina). Samples were run on an Illumina NovaSeq 6000 sequencer at the KUMC Genome Sequencing Facility. Reads from *.fastq files were mapped to the rat reference genome (Ensembl Rnor_5.0.78) using CLC Genomics Workbench 12.0 (Qiagen, Germantown, MD). Transcript abundance was expressed as reads per kilobase of transcript per million reads mapped (**RPKM**) and a *P* value of 0.05 was used as a cutoff for significant differential expression. Statistical significance was calculated by empirical analysis of digital gene expression followed by Bonferroni’s correction. Pathway analysis was performed using Metascape (22).

### Spermatocyte chromosome spreads and inmunostaining

Spermatocyte chromosome spreads and immunofluorescence staining were performed as previously reported (**23**) with modifications. Briefly, *Taf7l ^Xm+/Y^* and *Taf7l^Xm−/Y^*rat testes at PND 30 were dissected, tunica albuginea removed and deposited in Testis Incubation Medium (**TIM**) (104 mM NaCl, 45 mM KCl, 1.2 mM MgSO_4_, 0.6 mM KH_2_PO_4_, 6.0 mM sodium lactate, 1.0 mM sodium pyruvate, 0.1% glucose) with collagenase (2 mg/ml; C9891, Sigma) followed by an one h incubation with shaking at 550 revolutions per minute (**RPM**) at 32°C. After incubation, dissociated tubules were washed three times by centrifugation for 1 min at 600 RPM at room temperature. Separated tubules were resuspended in TIM with 200 μl of 0.25% trypsin (25200-056, Gibco Chemicals-ThermoFisher) and DNase I (4 μg/ml; DN-25, Sigma), incubated for 15 min at 32°C, and recovered by centrifugation at 550 RPM. Cells were passed through a 70-μm cell strainer, centrifuged for 5 min at 1200 RPM, resuspended with a DNAaseI solution (400 ug/ml), and washed three times with TIM. A 10 μL cell suspension was layered on 90 μL of 75 mM sucrose solution and incubated for 8 min at room temperature. Superfrost glass slides received 100 μL 1% paraformaldehyde (**PFA**) containing 0.15% Triton, pH 9.3. Cell suspensions (45 μL) was added, swirled three times, and dried in a closed slide box for three h, followed by drying with a half-open lid for two h at room temperature. Slides were then washed two times for three min in milli-Q water on a shaker, and one time for five min with 0.4% PhotoFlow, air-dried and stored in –80°C.

Slides containing spermatocyte spreads were blocked for 30 min at room temperature in 100 mL solution containing 2% BSA and 0.3% Tween-20 in PBS. Slides were incubated with primary antibody overnight in a humid chamber at 4°C. SCYP1 rabbit antibody (1:200; NB300-229, Novus, Centennial, CO), SYCP3 mouse antibody (1:200; SC-74569, Santa Cruz Biotechnology, Dallas, TX), and γH2AX rabbit antibody (1:500; 05-636, Millipore) were used in the analyses. Slides were washed three times for 10 min in PBS, then incubated with secondary antibody for 45 min at 37°C in a humid chamber. Alexa Fluor 568-conjugated goat anti-rabbit IgG (1:500; A11011, ThermoFisher) or Alexa Fluor 488-conjugated goat anti-mouse IgG (1:500; A11001, ThermoFisher) were used as secondary antibodies. Slides were washed three times for five min each and mounted with DAPI Fluoromount-G (0100-20, SouthernBiotech, Birmingham, AL).

### Statistical analysis

Student’s *t*-test, Welch’s *t*-test, Dunnett’s test, or Steel test were performed, where appropriate, to evaluate the significance of the experimental manipulations. Results were deemed statistically significant when *P*<0.05.

## RESULTS

### Expression of *Taf7l* in the rat

TAF7L is localized to cells at various stages of male germ cell differentiation in the mouse testis (11). The distribution of TAF7L in rat testis has not been examined. Initially we evaluated the expression of *Taf7l* transcripts during spermatogenesis in rat testis by *in situ* hybridization at postnatal day (**PND**) 10, 30 and 60 (**Fig. 1**). *Taf7l* transcripts were prominently localized to preleptotene spermatocytes, where they exhibited co-localization with *Stra8*, a known regulator of germ cell development and meiosis (24–27). These findings place TAF7L in position to potentially affect meiotic prophase I during spermatogenesis.

**Figure 1.**
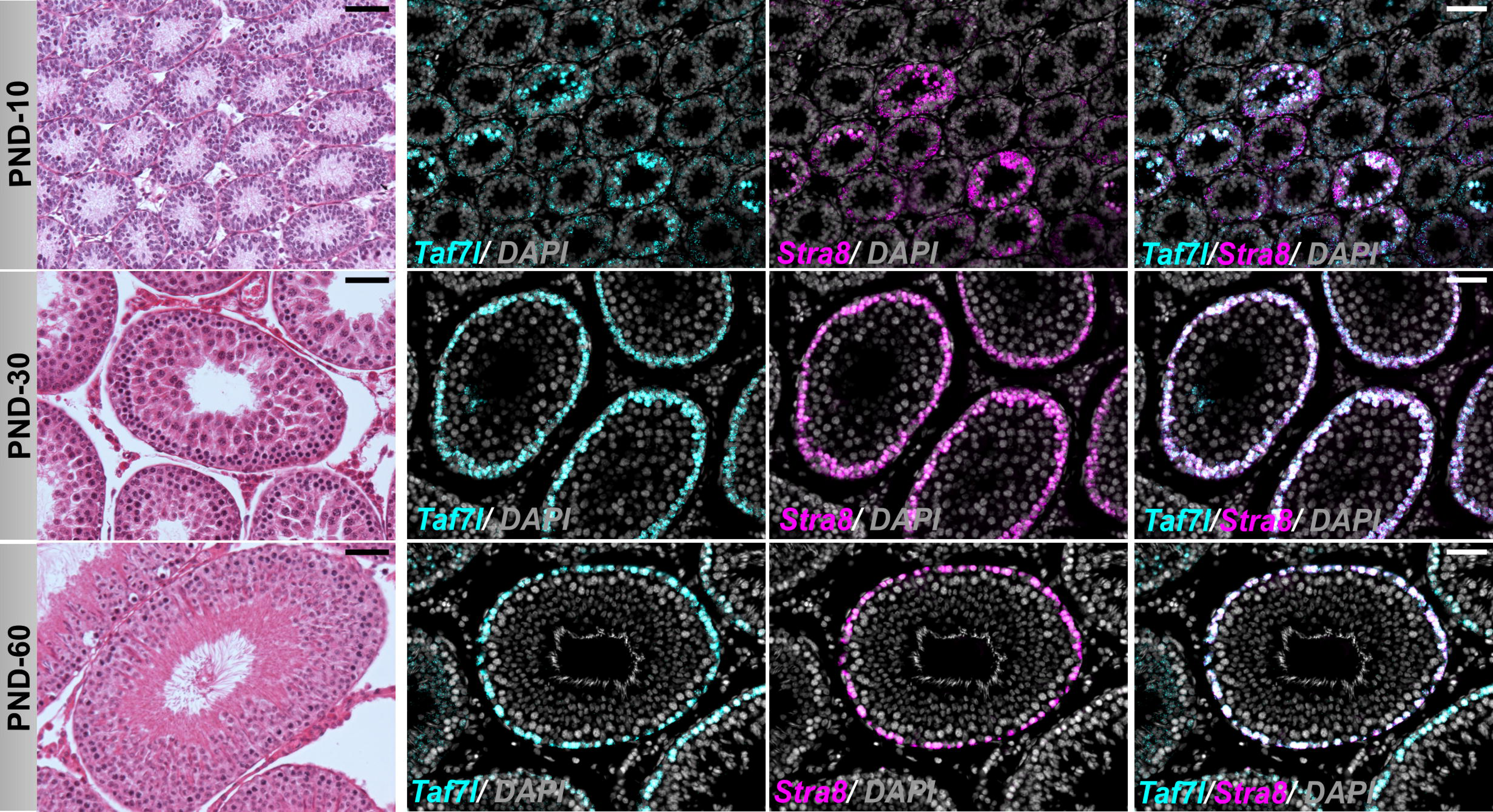
*Taf7l* transcript localization in rat testes. Left panels, representative histological sections of testes at postnatal day (**PND**) 10, 30, and 60 stained with hematoxylin and eosin. Representative in situ hybridization for *TAf7l* and *Stra8* in PND 10, 30, and 60 sections of testes (**right**). Scale bar, 50 μm.

### Generation of an *Taf7l* mutant rat model

We examined the role of TAF7L in regulating spermatogenesis in the rat using CRISPR/Cas9 genome editing. A mutant rat model possessing a 110 bp deletion within the *Taf7l* gene was generated (**Fig. 2**). The deletion included part of Exon 3 and led to a frameshift and premature stop codon. The deletion effectively removed the TAFII55 protein conserved region of TAF7L (**Fig. 2A-B**). The *Taf7l* mutation was successfully transmitted through the germline. A rat colony possessing the *Taf7l* mutation was established and maintained via heterozygous female x wild type male breeding which produced the predicted Mendelian ratio. Wild type (*Taf7l^Xm+^*) and mutant (*Taf7l^Xm-^*) polymerase chain reaction (**PCR**) products (wild type allele: 550 bp versus mutant allele: 440 bp) could be readily distinguished for genotyping (**Fig. 2C**). The *Taf7l* gene was successfully disrupted in the rat.

**Figure 2.**
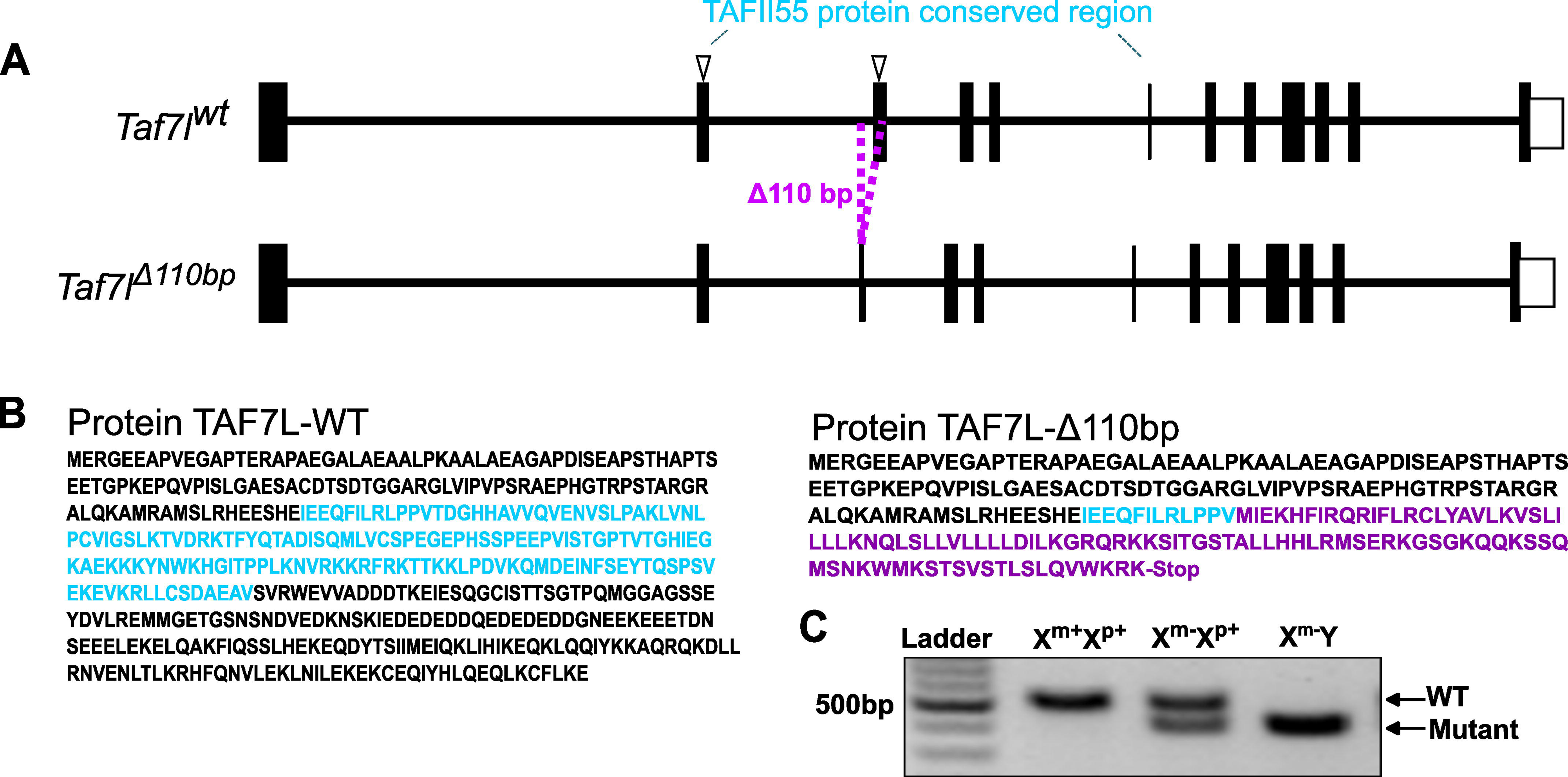
In vivo genome editing of the rat *Taf7l* locus. **A**) Schematic representation of the rat *Taf7l* gene (*Taf7l^wt^*, NM_001135877) and the mutant *Taf7l* allele with a 110 bp deletion (*Taf7l*^Δ*110bp*^). Arrowheads at Exons 2 and 3 correspond to the 5’ and 3’ guide RNAs used in the genome editing. **B**) Amino acid sequences for predicted the wild type (**TAF7L-WT**) and mutant (**TAF7L-**Δ**110bp**) TAF7L proteins. The magenta sequence corresponds to the frameshift in Exon 3 and the premature stop codon. The highlighted amino acid sequence in light blue corresponds to the TAFII55 protein conserved region. **C**) Wild type (X^m+^X^p+^ or X^m+^Y), heterozygous females (X^m-^/X^p+^), and homozygous mutant males (X^m-^/Y) genotypes were determined by PCR.

### Small testes and sterility in the *Taf7l^Xm-/Y^* rat

*Taf7l^Xm−/Xp+^* females presented as healthy and produced healthy offspring. In contrast, *Taf7l^Xm-/Y^* males were infertile (**Fig. 3A**). Although mutant males maintained normal body weight, their testes were significantly smaller than littermate controls (*Taf7l^Xm+/Y^*) at PND 30 and 60 (**Fig. 3B, Supplementary** Fig. 1). Epididymal sperm were not present in *Taf7l^Xm-/Y^* rats (**Fig. 3C**).

**Figure 3.**
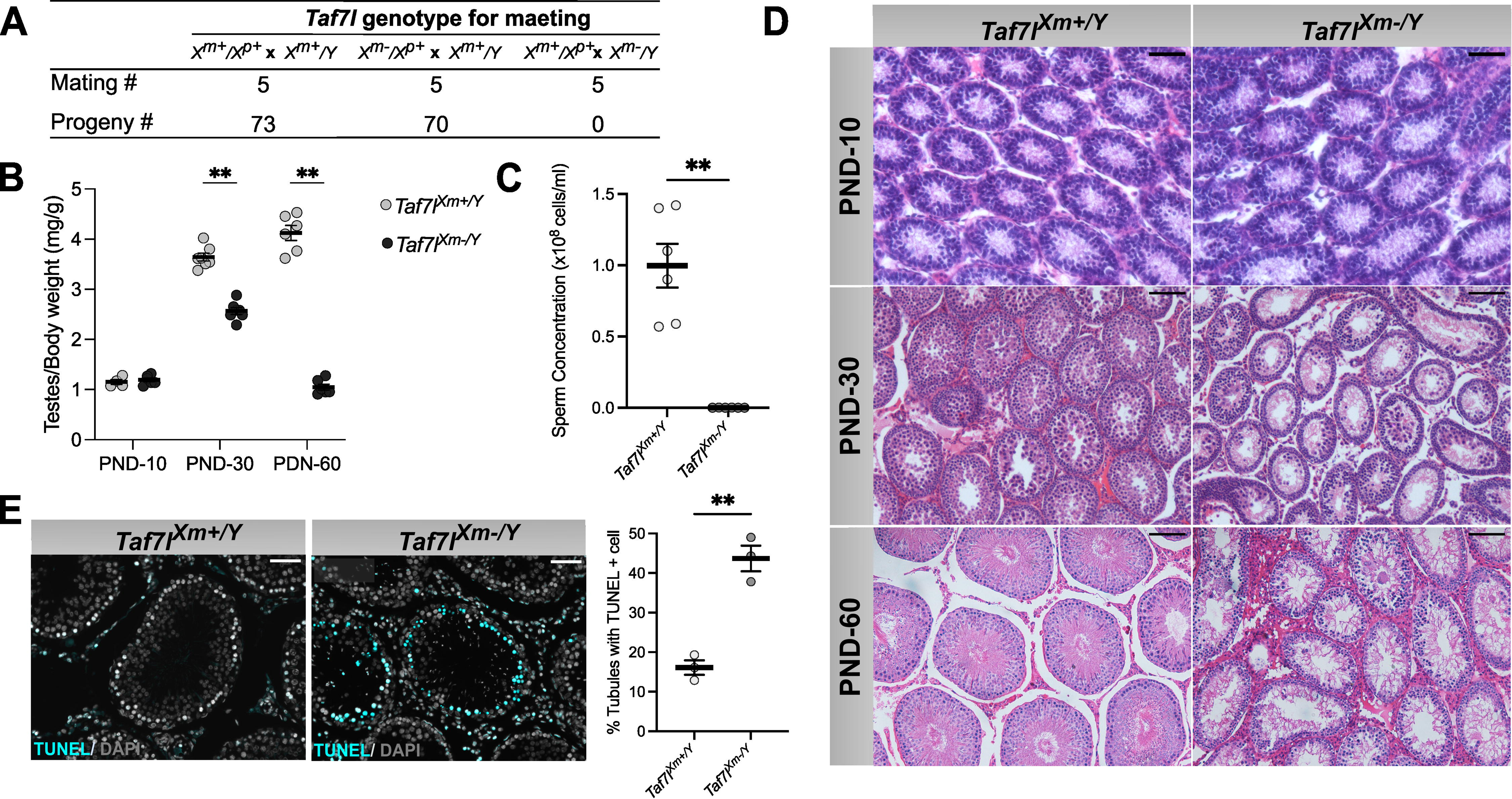
*Taf7l* mutation impairs testis development and function in the rat. **A**) Progeny produced from mating wild type (**WT**) males (*Taf7l^Xm+/Y^*) and WT females (*Taf7l^Xm+/Xp+^*), heterozygous females (*Taf7l^Xm-/Xp+^*) and WT males (*Taf7l^Xm+/Y^*), and WT females (*Taf7l^Xm+/Xp+^*) and homozygous mutant males (*Taf7l^Xm-/Y^*). **B**) Weights of testes related to body weight obtained from *Taf7l^Xm+/Y^* or *Taf7l^Xm-/Y^* rats at postnatal day (**PND**) 10, 30 and 60. Data are presented as the mean ± SEM. Dots represent biological replicates per condition \*\**p<0.005*. **C**) Sperm quantification for *Taf7l^Xm+/Y^* and *Taf7l^Xm-/Y^*rats at PND 60. Data are presented as the mean ± SEM. Dots represent biological replicates per condition (n=6); unpaired *t* tests were used to assess for the significance of differences between the groups, \*\**p<0.005*. **D**) Hematoxylin and eosin staining of *Taf7l^Xm+/Y^* and *Taf7l^Xm-/Y^* testes at PND 10, 30 and 60. Scale bar PND 10 = 50 μm, PND 30 and 60 = 100 μm. **E**) TUNEL staining of PND 60 *Taf7l^Xm+/Y^* or *Taf7l^Xm-/Y^* testes. Data are presented as the mean ± SEM. Dots represent biological replicates per condition. Unpaired *t* tests were used to assess for the significance of differences between the groups, \*\**p<0.005*. Scale bars = 50 μm.

Histologic analyses were performed on testes from PND 10, 30, and 60 *Taf7l^Xm+/Y^* and *Taf7l^Xm-/Y^* animals. TAF7L-dependent histological differences were not evident in testes from PND10 (**Fig. 3D, top panel**) but were apparent on PND 30 and 60 (**Fig. 3D, middle and bottom panels**). Control PND 30 and 60 testes (*Taf7l^Xm+/Y^*) had a full array of spermatogenic cells, including spermatogonia, spermatocytes, and round and elongated spermatids, and possessed spermatocytes in the seminiferous tubules, whereas TAF7L mutant PND 30 and 60 mutant testes (*Taf7l^Xm-/Y^*) exhibited spermatogenic arrest with an absence of spermatids (**Fig. 3D, middle and bottom panels**).

To further characterize wild type and *Taf7l^Xm-/Y^* testes, we performed terminal dUTP nick-end labeling (**TUNEL**) analyses. Seminiferous tubules from *Taf7l^Xm-/Y^* mutant testes possessed significantly more TUNEL-positive cells than did control testes (**Fig. 3E**).

In summary, TAF7L deficiency results in small testes, an arrest in spermatogenesis, and an increase in the appearance of apoptotic cells within the seminiferous tubules.

### TAF7L is required for spermatogenesis stage specific gene expression in the rat

To obtain additional insight into the involvement TAF7L in spermatogenesis we performed bulk RNA-sequencing (**RNA-seq**) on PND 60 *Taf7l^Xm+/Y^*(n=4) and *Taf7l^Xm-/Y^* (n=4). testes. RNA-seq analyses yielded 10,065 differentially expressed genes (**DEGs**), which included 5,559 downregulated and 4,506 upregulated genes in *Taf7l^Xm-/Y^* testes (**Fig. 4A**). Pathway analysis of the top 3,000 DEGs highlighted gene involvement in regulating male gonad development, spermatogenesis, meiotic cell cycle, and meiosis (**Fig. 4B**).

**Figure 4.**
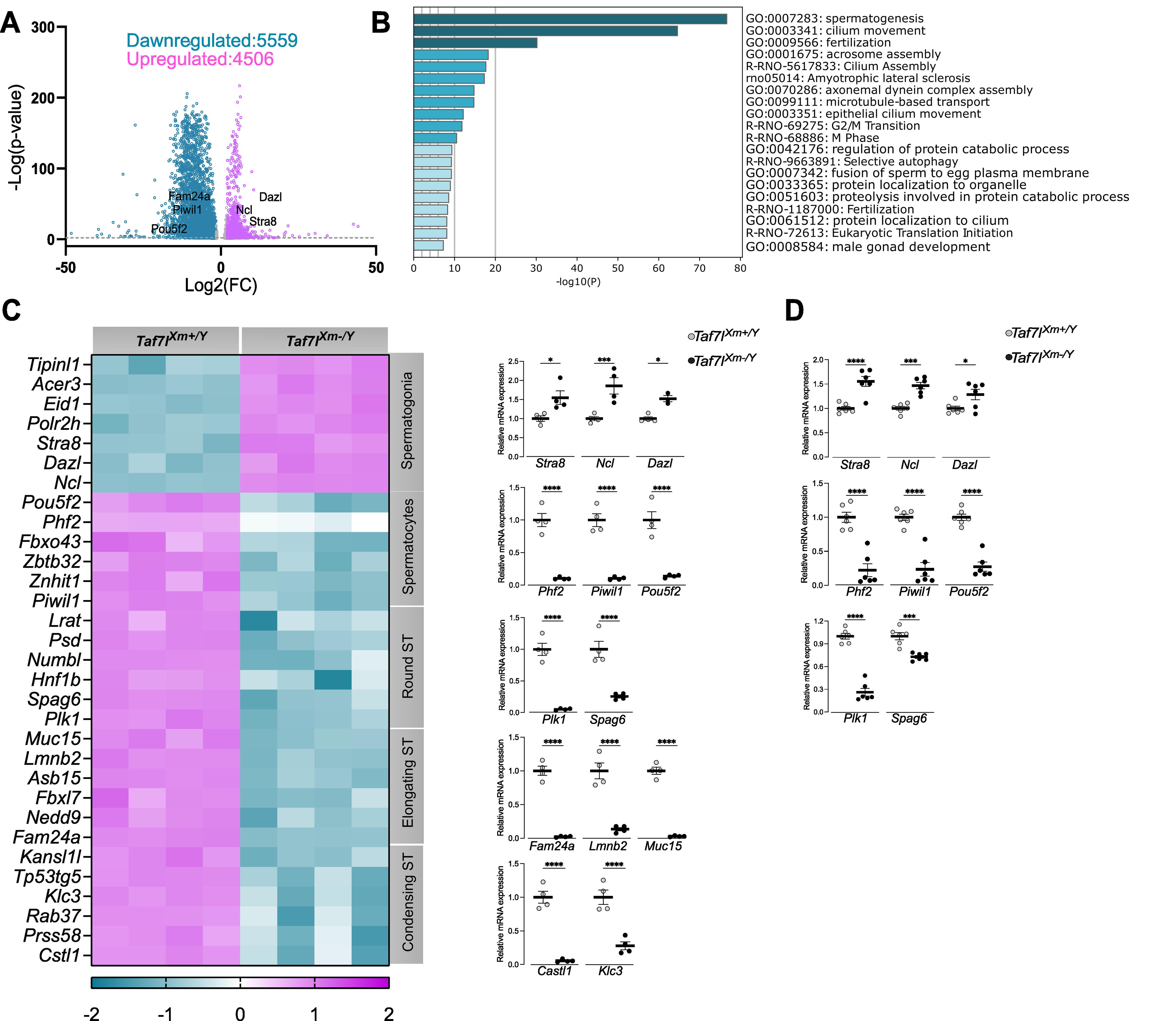
TAF7L is required for spermatogenesis stage specific gene expression in the rat as assessed by RNA-sequencing (RNA-seq). **A**) Volcano plot of differentially expressed genes (**DEGs**) in wild type (*Taf7l^Xm+/Y^*) and mutant (*Taf7l^Xm-/Y^*) testes at postnatal day (**PND**) 60 (n=4). Cyan dots represent significantly down-regulated transcripts with a logarithm to base two-fold change of less than or equal to −2. Magenta dots represent significantly up-regulated transcripts with a logarithm to base two-fold change of greater than or equal to 2. The dotted line represents p≤0.05. **B**) Gene ontology analysis of downregulated DEGs. **C**) Heat map depicting selected DEGs for *Taf7l^Xm+/Y^* and *Taf7l^Xm-/Y^* PND-60 testes (**left**). The heat map also highlights genes expressed by specific spermatogenic cell types, which were determined by integrating the bulk RNA-seq data with published single cell RNA-seq datasets for the rat (**Guan et al., 2022**). ST, spermatid. The heat map color key represents z-scores of reads per kilobase per million mapped reads. Validation of RNA-seq results by RT-qPCR for each group at PND-60 is shown on the **right**; ST, spermatid**. D**) Validation of RNA-seq results by RT-qPCR at PND-30. *Gapdh* mRNA expression levels were used as a control for normalization. In all graphs, data are presented as the mean ± SEM. Dots represent biological replicates per condition (n=4-6). Unpaired *t* tests were used to assess for the significance of differences between the groups, \**p<0.05,***p<0.001, ****p<0.0001*.

Next, we integrated published single cell RNA-seq (**scRNA-seq**) datasets of adult rat testicular cells (28) with our bulk RNA-seq datasets from PND 60 *Taf7l^Xm+/Y^* and *Taf7l^Xm-/Y^*testes to identify specific transcript-associations with each spermatogenic cell type. The dataset integration indicated that TAF7L deficiency led to an enrichment of spermatogonia genes, and a downregulation of genes associated to spermatocytes and spermatids (**Fig. 4C**). These findings were validated by reverse transcriptase-quantitative polymerase chain reaction (**RT-qPCR**) on PND 30 and 60 *Taf7l^Xm+/Y^* and *Taf7l^Xm-/Y^* testes as well as by *in situ* hybridization (**Fig. 4C-D, Supplementary** Fig. 2). The findings indicate that in the rat testis TAF7L disruption leads to meiotic arrest.

### TAF7L deficiency leads to an arrest in spermatogenesis at the late zygonema stage

To understand the involvement of TAF7L in meiosis we evaluated synapsis on spermatocyte spreads by immunostaining for SYCP1, a central element of the synaptonemal complex, and SYCP3, a lateral element of the synaptonemal complex, from PND 30 *Taf7l^Xm+/Y^* and *Taf7l^Xm-/Y^* testes. Meiotic leptonema and zygonema stages were present in wild type and TAF7L deficient specimens; however, only wild type spermatocytes progressed to the pachynema stage (**Fig. 5A**). We next examined the phosphorylation of H2A histone family member X (**H2AX**), which is formed in response to double-strand DNA breaks and disappears after synapsis of autosomal chromatin during the pachynema stage and subsequently, forms on the X and Y chromosomes in the sex body (29, 30). In *Taf7l^Xm+/Y^*most of the phosphorylated H2AX (γ**H2AX**) signal disappeared as autosomes synapsed, leaving brightly stained sex bodies in pachynema (**Fig. 5B**). On the other hand, most cells from *Taf7l^Xm-/Y^* testes do not possess sex bodies (Fig. 5B). Synapsis was not complete in these cells (**Fig. 5B**). Most of the cells in the *Taf7l^Xm-/Y^* testis were arrested in zygonema with the γH2AX signal present on autosomal chromatin (**Fig. 5B**). The percentage of cells in pachynema with an γH2AX signal on sex bodies was significantly decreased in *Taf7l^Xm-/Y^* (**Fig. 5C**).

**Figure 5.**
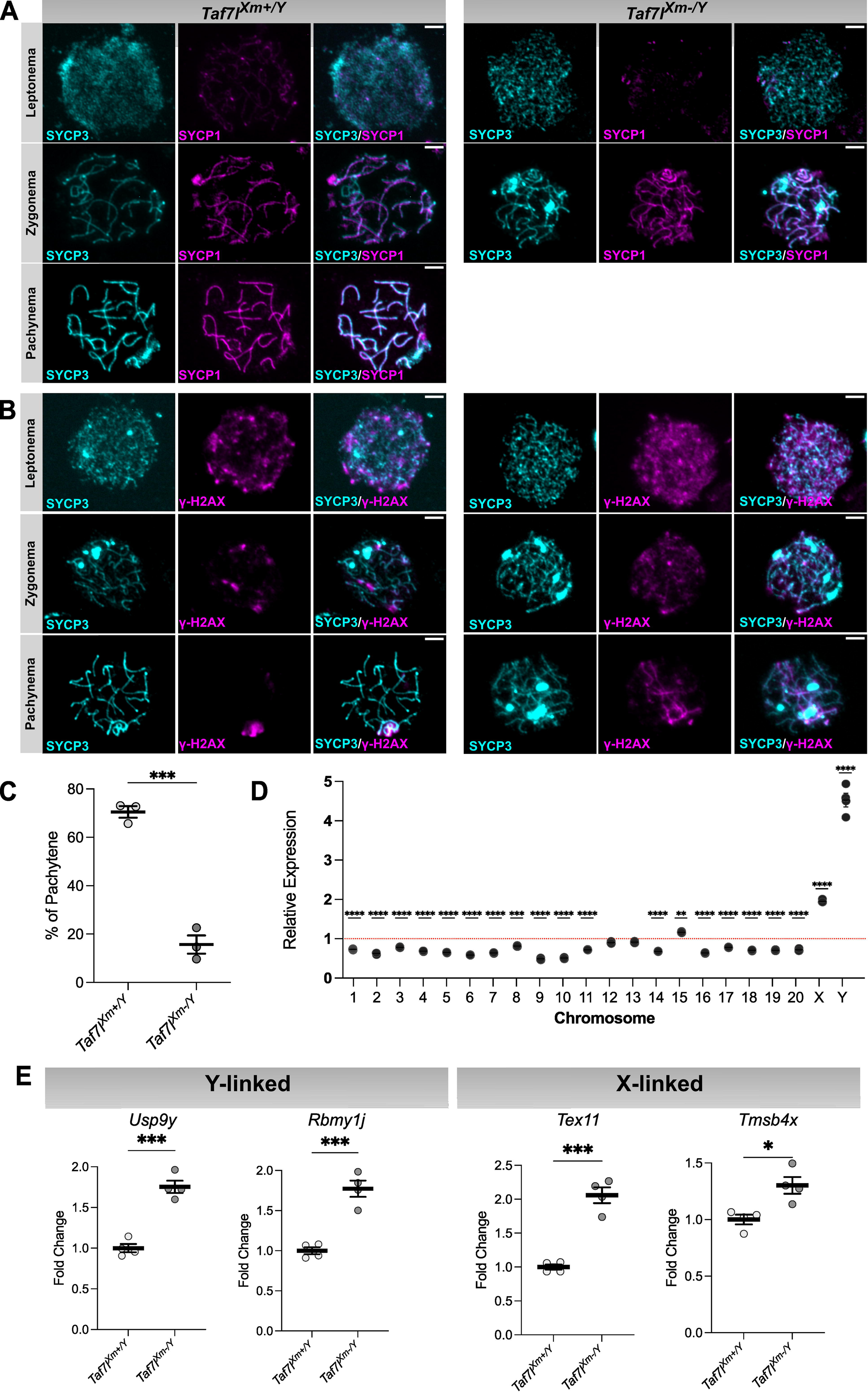
Meiotic arrest in *Taf7l* mutant rat. **A)** Meiotic progression in wild type (*Taf7l^Xm+/Y^*) and *Taf7l* mutant (Taf7l^Xm-/Y^) spermatocytes at postnatal day (**PND**) 30. Representative immunostaining for SYCP3 and SYCP1 in chromosome spreads are shown. Scale bar = 5 μm. **B**) Representative immunostaing for SYCP3 and γH2AX in chromosome spreads are shown. **C**) Percentage of cell in pachynema stage (presence of sex body) in *Taf7l^Xm+/Y^* and *Taf7l^Xm-/Y^* testes at PND-30. Data are presented as the mean ± SEM. Dots represent biological replicates per condition (n=3); unpaired *t* test, *\*\*\*p<0.001.* **D**) Relative levels of expression per chromosome for *Taf7l* mutant averaged across all genes expressed (p<0.05) in testes at PND-60 determined by RNA-seq analysis. Unpaired *t* test, \*\**p*<0.01, \*\*\**p*<0.001, *\*\*\*\*p<0.0001*. Red dotted line represents *Taf7l^Xm+/Y^* relative expression. **E**) Relative expression levels of representative Y-linked (*Usp9y* and *Rbmy1j*) and X-linked (*Tex11* and *Tmsb4x*) genes from *Taf7l^Xm+/Y^* and *Taf7l^Xm-/Y^*testes at PND-30 determined by RT-qPCR. *Gapdh* mRNA expression was used as a control. Data are presented as the mean ± SEM. Dots represent biological replicates per condition (n=4); unpaired *t* test, \**p<0.05,***p<0.001*.

### Failure to undergo MSCI in *Taf7l^Xm-/Y^* testes

The condensation of the X and Y chromosomes to form the sex body contributes to MSCI (31, 32). We next determined whether the failure of sex body formation in *Taf7l^Xm-/Y^*males resulted in a disruption of MSCI and compared the expression of all dysregulated genes from our RNA-seq data of PND 60 *Taf7l^Xm+/Y^*and *Taf7l^Xm-/Y^* testis. We found that most autosomal genes were downregulated in the *Taf7l^Xm-/Y^* (**Fig. 5D**). In contrast, X-linked genes and Y-linked genes were overexpressed in *Taf7l^Xm-/Y^* testis (**Fig. 5D**). We validated two X-linked genes (*Tex11*, *Tmsb4x*) and two Y-linked genes (*Rbmy1j*, *Usp9y*) by RT-qPCR in PND 30 *Taf7l^Xm+/Y^* and *Taf7l^Xm-/Y^* testes where the population of spermatocytes is more homogenous (**Fig. 5E**). These results indicate that MSCI is impaired in the *Taf7l^Xm-/Y^* testis.

## DISCUSSION

In this study, we evaluated the biology of TAF7L in male reproduction. A global mutant rat model was used to assess the role of TAF7L in the rat testis. Disruption of *Taf7l* resulted in a failure of spermatogenesis progression, secondary to an arrest in meiosis, failure of sperm production, and male infertility. TAF7L contributed to the regulation of MSCI. The TAF7L mutant phenotype was robust and offers a useful model for investigating the gene regulatory network controlling spermatogenesis in the testis.

TAF7L deficiency led to an arrest in spermatogenesis at the late zygonema stage and a failure of MSCI. These events are precisely regulated. TAF7L likely coordinates transcriptional events pivotal to the progression of these key stages of spermatogenesis. Previous reports have highlighted the role of sex chromosome genes in spermatogenesis (33, 34). Genes predominantly expressed in spermatogonia and Sertoli cells independently accumulated on X chromosomes across mammals (34). During the pachytene stage of meiosis in male mammals, X and Y chromosomes are transcriptionally silenced by MSCI (35–37). MSCI is conserved in mammals (34) and is essential for normal male fertility. In this study, we show that TAF7L deficiency disrupted XY body formation, leading to defective MSCI. These results reveal a novel role of TAF7L in the regulation of male meiotic prophase and provides a potential regulatory mechanism for the MSCI process. The mechanism of TAF7L action on MSCI remains undetermined.

In the rat, TAF7L plays an essential role in spermatogenesis. *Taf7l* is expressed prominently in spermatogonia where it co-localizes with *Stra8*. These observations are consistent with single cell-RNA-sequencing and spatial transcriptomics data generated for the mouse and human (38–40). Although, independent validation of TAF7L expression in the human testis would is needed, it appears that TAF7L may be contributing to similar biology within the testes of the rat, mouse and human. However, testis phenotypes following *Taf7l* gene disruption in the rat and mouse are different. Spermatogenesis is arrested in the *Taf7l* null rat resulting in infertility, whereas *Taf7l* null mice possess disturbances in the production of healthy sperm and exhibit subfertility (12, 13). The hypomorphic mouse testis phenotype is suggestive of some compensation from another TAF family member, possibly TAF7. These observations highlight differences in the biology of TAF7L in the mouse, rat, and human testis and the need for more in depth analysis of TAF7L as a regulator of spermatogenesis and identification of compensatory pathways replacing TAF7L in the mouse and human.

## ACKNOWLEDGEMENTS

The research was supported by postdoctoral fellowships from the Kansas Idea Network of Biomedical Research Excellence, P20 GM103418 (A.M-I., E.D), Lalor Foundation (A.M-I., ED), and NIH grants (MJS:HD020676, HD099638, HD105734; NW:HD103888) and the Sosland Foundation. We thank Dr. Saher Sue Hammoud, Dr. Richard N. Freiman and Dr. Haiqi Chen for helpful discussions at early stages of the project. We also thank Stacy Oxley and Brandi Miller for administrative assistance.

## AUTHOR CONTRIBUTIONS

A.M.I., X.Z, N.W and M.J.S. conceived and designed the research; A.M.I., E.D., K.I. performed experiments; A.M.I, N.W and M.J.S. analyzed the data and interpreted results of experiments; A.M.I., and M.J.S. prepared figures and manuscript; All authors read, contributed to editing, and approved the final version of manuscript.

## DATA AVAILABILITY

RNA-seq datasets are available at the Gene Expression Omnibus (**GEO**) database, https://www.ncbi.nlm.nih.gov/geo/ (in progress). All data generated and analyzed in this study are included in the published article and supporting files. Resources generated from the research are available from the corresponding author upon reasonable request.

## REFERENCES

1. P. Sassone-Corsi, Transcriptional checkpoints determining the fate of male germ cells. Cell 88, 163–166 (1997).

2. S. Kimmins, N. Kotaja, I. Davidson, P. Sassone-Corsi, Testis-specific transcription mechanisms promoting male germ-cell differentiation. Reprod. Camb. Engl. 128, 5–12 (2004).

3. J. M. Oatley, R. L. Brinster, Regulation of spermatogonial stem cell self-renewal in mammals. Annu. Rev. Cell Dev. Biol. 24, 263–286 (2008).

4. G. J. Veenstra, A. P. Wolffe, Gene-selective developmental roles of general transcription factors. Trends Biochem. Sci. 26, 665–671 (2001).

5. A. Hochheimer, R. Tjian, Diversified transcription initiation complexes expand promoter selectivity and tissue-specific gene expression. Genes Dev. 17, 1309– 1320 (2003).

6. M. A. Hiller, T. Y. Lin, C. Wood, M. T. Fuller, Developmental regulation of transcription by a tissue-specific TAF homolog. Genes Dev. 15, 1021–1030 (2001).

7. M. Hiller, et al., Testis-specific TAF homologs collaborate to control a tissue-specific transcription program. Dev. Camb. Engl. 131, 5297–5308 (2004).

8. P. J. Wang, J. R. McCarrey, F. Yang, D. C. Page, An abundance of X-linked genes expressed in spermatogonia. Nat. Genet. 27, 422–426 (2001).

9. P. J. Wang, D. C. Page, Functional substitution for TAF(II)250 by a retroposed homolog that is expressed in human spermatogenesis. Hum. Mol. Genet. 11, 2341–2346 (2002).

10. A. E. Falender, et al., Maintenance of spermatogenesis requires TAF4b, a gonad-specific subunit of TFIID. Genes Dev. 19, 794–803 (2005).

11. J.-C. Pointud, et al., The intracellular localisation of TAF7L, a paralogue of transcription factor TFIID subunit TAF7, is developmentally regulated during male germ-cell differentiation. J. Cell Sci. 116, 1847–1858 (2003).

12. Y. Cheng, et al., Abnormal sperm in mice lacking the Taf7l gene. Mol. Cell. Biol. 27, 2582–2589 (2007).

13. H. Zhou, et al., Taf7l cooperates with Trf2 to regulate spermiogenesis. Proc. Natl. Acad. Sci. U. S. A. 110, 16886–16891 (2013).

14. K. Stouffs, et al., The role of the testis-specific gene hTAF7L in the aetiology of male infertility. Mol. Hum. Reprod. 12, 263–267 (2006).

15. O. Akinloye, J. Gromoll, C. Callies, E. Nieschlag, M. Simoni, Mutation analysis of the X-chromosome linked, testis-specific TAF7L gene in spermatogenic failure. Andrologia 39, 190–195 (2007).

16. M. Heidari, M. Sheidai, Z. Noormohammadi, G. A. Roozbahani, N. Kalhor, Association study of TAF7L two SNPs with oligozoospermia patients. Gene Rep. 9, 127–130 (2017).

17. L. Ling, et al., Genetic characterization of a missense mutation in the X-linked TAF7L gene identified in an oligozoospermic man†. Biol. Reprod. 107, 157–167 (2022).

18. H. Bai, et al., Deleterious variants in TAF7L cause human oligoasthenoteratozoospermia and its impairing histone to protamine exchange inducing reduced in vitro fertilization. Front. Endocrinol. 13, 1099270 (2022).

19. T. Kaneko, Genome Editing of Rat. Methods Mol. Biol. Clifton NJ 1630, 101–108 (2017).

20. K. Iqbal, P. Dhakal, S. H. Pierce, M. J. Soares, Catechol-O-methyltransferase and Pregnancy Outcome: an Appraisal in Rat. Reprod. Sci. Thousand Oaks Calif 28, 462–469 (2021).

21. H. Ecroyd, K. L. Asquith, R. C. Jones, R. J. Aitken, The development of signal transduction pathways during epididymal maturation is calcium dependent. Dev. Biol. 268, 53–63 (2004).

22. Y. Zhou, et al., Metascape provides a biologist-oriented resource for the analysis of systems-level datasets. Nat. Commun. 10, 1523 (2019).

23. M. Boekhout, et al., REC114 Partner ANKRD31 Controls Number, Timing, and Location of Meiotic DNA Breaks. Mol. Cell 74, 1053–1068.e8 (2019).

24. A. E. Baltus, et al., In germ cells of mouse embryonic ovaries, the decision to enter meiosis precedes premeiotic DNA replication. Nat. Genet. 38, 1430–1434 (2006).

25. J. Koubova, et al., Retinoic acid regulates sex-specific timing of meiotic initiation in mice. Proc. Natl. Acad. Sci. U. S. A. 103, 2474–2479 (2006).

26. E. L. Anderson, et al., Stra8 and its inducer, retinoic acid, regulate meiotic initiation in both spermatogenesis and oogenesis in mice. Proc. Natl. Acad. Sci. U. S. A. 105, 14976–14980 (2008).

27. Q. Zhou, et al., Expression of stimulated by retinoic acid gene 8 (Stra8) in spermatogenic cells induced by retinoic acid: an in vivo study in vitamin A-sufficient postnatal murine testes. Biol. Reprod. 79, 35–42 (2008).

28. X. Guan, et al., Single-cell RNA sequencing of adult rat testes after Leydig cell elimination and restoration. Sci. Data 9, 106 (2022).

29. S. K. Mahadevaiah, et al., Recombinational DNA double-strand breaks in mice precede synapsis. Nat. Genet. 27, 271–276 (2001).

30. M. Barchi, et al., Surveillance of different recombination defects in mouse spermatocytes yields distinct responses despite elimination at an identical developmental stage. Mol. Cell. Biol. 25, 7203–7215 (2005).

31. J. M. A. Turner, Meiotic Silencing in Mammals. Annu. Rev. Genet. 49, 395–412 (2015).

32. K. G. Alavattam, S. Maezawa, P. R. Andreassen, S. H. Namekawa, Meiotic sex chromosome inactivation and the XY body: a phase separation hypothesis. Cell. Mol. Life Sci. CMLS 79, 18 (2021).

33. J. L. Mueller, et al., The mouse X chromosome is enriched for multicopy testis genes showing postmeiotic expression. Nat. Genet. 40, 794–799 (2008).

34. F. Murat, et al., The molecular evolution of spermatogenesis across mammals. Nature 613, 308–316 (2023).

35. B. D. McKee, M. A. Handel, Sex chromosomes, recombination, and chromatin conformation. Chromosoma 102, 71–80 (1993).

36. W. Yan, J. R. McCarrey, Sex chromosome inactivation in the male. Epigenetics 4, 452–456 (2009).

37. J. M. A. Turner, Meiotic sex chromosome inactivation. Dev. Camb. Engl. 134, 1823–1831 (2007).

38. A. N. Shami, et al., Single-Cell RNA Sequencing of Human, Macaque, and Mouse Testes Uncovers Conserved and Divergent Features of Mammalian Spermatogenesis. Dev. Cell 54, 529–547.e12 (2020).

39. J. Guo, et al., Single-cell analysis of the developing human testis reveals somatic niche cell specification and fetal germline stem cell establishment. Cell Stem Cell 28, 764–778.e4 (2021).

40. S. Rajachandran, et al., Dissecting the spermatogonial stem cell niche using spatial transcriptomics. Cell Rep. 42, 112737 (2023).

