## Supplementary Material for "TAF7L REGULATES EARLY STAGES OF MALE GERM CELL DEVELOPMENT"

**This PDF file includes:**

Figures S1 to S2

Tables S1

**Other supporting materials for this manuscript include the following:**

Datasets S1 to S2

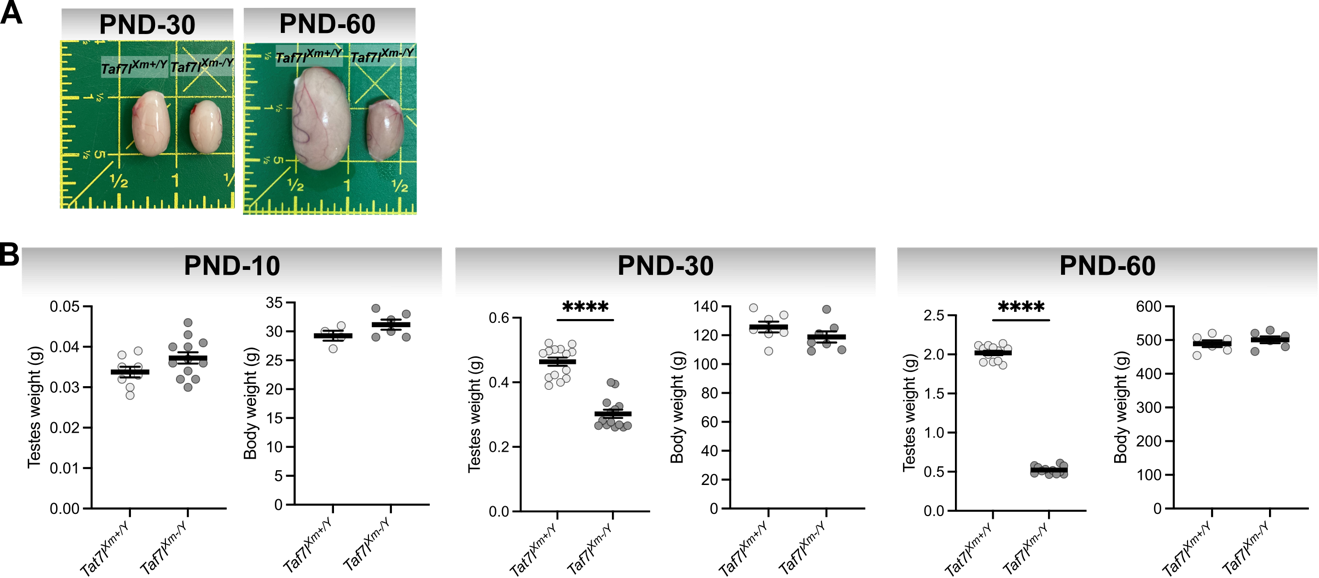

Fig. S1. *Taf7l* mutation impairs testis development in the rat. A) Testes from *Taf7l^Xm+/Y^* and *Taf7l^Xm-/Y^* males at postnatal day (PND) 30 and 60. B) Testes weights and body weights obtained from *Taf7l^Xm+/Y^* or *Taf7l^Xm-/Y^* males at PND 10, 30 and 60. Data are presented as the mean ± SEM. Dots represent biological replicates per condition (n=4-6); unpaired *t* test*****p<0.0005*.

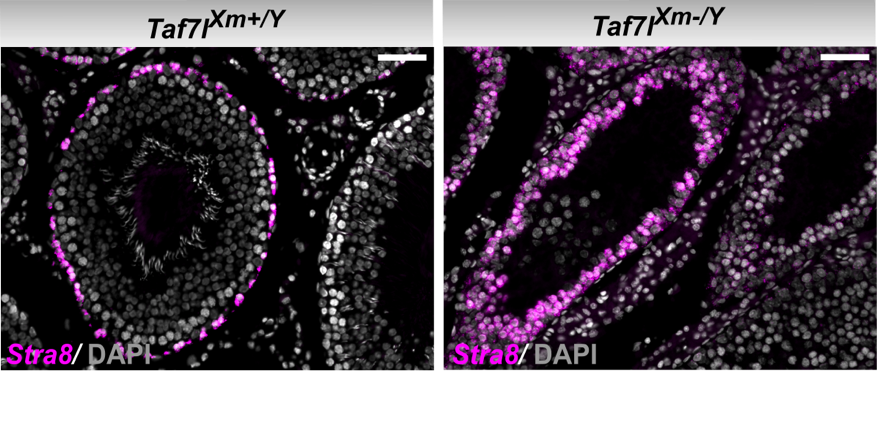

Fig. S2. Disruption of *Taf7l* results in an expansion of spermatogonial cells. Representative *Stra8* *in situ* hybridization images of postnatal day 60 testes sections from *Taf7l^Xm+/Y^* (left panel) and *Taf7l^Xm-/Y^* (right panel) males. Scale bar = 50mm.

Table S1. Primers used for RT-qPCR.

| **Gene** | **Forward primer** | **Reverse primer** |
| --- | --- | --- |
| *Pou5f2* | CAGCCAGGCAGCCACAAGCAGG | AGCTCCTTGGCCAGCTGCTCC |
| *Piwil1* | GCGACCCGGCGTGAGGAGCC | GGTCCAGCAGCGTCTGGTCC |
| *Phf2* | CGGACGACGGGGAGCTGAAG | TGGGGATGAGGAGAGTCTCC |
| *Znhit1* | ACGCCGGCCTCCCCCAGCTGG | GGGCCCTCAGATGCACTCAGG |
| *Spag6* | GGGGCCCTGGACTCACTGGTG | ATCACTGAGGGCCGAGGCAGC |
| *Plk1* | GCTCAAGCCCCATCAGAAGGA | GGCTCGGTCAGCGCCTTCCTCC |
| *Lmnb2* | GGCCGCAGCCCTCGGTGACAA | TCCTGCAGACTCTGGCAGTGG |
| *Fam24a* | CAGTGAGGCCCATCGGCCCAT | CAGGGCCTTGGATATTTTCAGG |
| *Muc15* | ATTCCTCCCTTGGTTCAGGGC | CGGGTGTGGTCTTTGGAGGAG |
| *Cstl1* | CTGTGACTGGTGGGAGGCAG | TGTTGCCAGTTTGGGAGTGC |
| *Klc3* | GGCCATTGAGCTGGGACTGGG | CCTCACTGGCCCGCAGTCGCC |
| *Prss53* | CCACAGCCCCCAAGCTAAGAG | CACGCTGAGCTGCTTGAAGAC |
| *Usp26* | CCACAACCAATCTAGGTCCG | GTCACCGTGCTGTCCTGTTTG |
| *Tmsb4x* | CCTTCCAGCAACCATGTCTG | AATGGGCGAGGGCGAACAGC |
| *Usp9y* | GCCTCAAGAATTCCAGGACA | GCCATTCGCTCAGCGGTTAG |
| *Tex11* | GCTGTGACAGAGCCCTGAACC | CTAAGGTCAGCAATGCCATCC |
| *Rbmy* | GGAGGCCTCAACTTAGAGACC | GCATTCTTAGCATCTCCAGGAC |
| *Uba1y* | GTCCAGCTCGGTCCTGTCCA | AGCTGGCGGGAGTAAAGGCT |
